# Two Novel α-L-Arabinofuranosidases from *Bifidobacterium longum* subsp. *longum* belonging to Glycoside Hydrolase Family 43 Cooperatively Degrade Arabinan

**DOI:** 10.1101/379495

**Authors:** Masahiro Komeno, Honoka Hayamizu, Kiyotaka Fujita, Hisashi Ashida

## Abstract

Arabinose-containing poly-or oligosaccharides are suitable carbohydrate sources for *Bifidobacterium longum* subsp. *longum*, though their degradation pathways are poorly understood. In this study, we found that the gene expression levels of *bllj 1852* and *bllj 1853* from *B. longum* subsp. *longum* JCM 1217 were enhanced in the presence of arabinan. Both genes encode previously uncharacterized glycoside hydrolase (GH) family 43 enzymes. Subsequently, we cloned those genes and characterized the recombinant enzymes expressed in *Escherichia coli.* Both enzymes exhibited α-L-arabinofuranosidase activity toward synthetic *p*-nitrophenyl glycoside, but the specificities for L-arabinofuranosyl linkages were different. BLLJ_1852 catalyzed the hydrolysis of α1,2- and α1,3-L-arabinofuranosyl linkages found in the side chains of arabinan and arabinoxylan. BLLJ_1852 released L-arabinose 100 times faster from arabinan than from arabinoxylan but did not act on arabinogalactan. BLLJ_1853 catalyzed the hydrolysis of α1,5-L-arabinofuranosyl linkages found on the arabinan backbone. BLLJ_1853 released L-arabinose from arabinan but not from arabinoxylan or arabinogalactan. Both enzyme activities were largely suppressed with EDTA treatment, suggesting that they require divalent metal ions. BLLJ_1852 was moderately activated in the presence of all divalent cations tested, whereas BLLJ_1853 activity was inhibited by Cu^2+^. The GH43 domains of BLLJ_1852 and BLLJ_1853 are classified into GH43 subfamilies 27 and 22, respectively, but hardly share similarity with other biochemically characterized members in the corresponding subfamilies.

**IMPORTANCE:** We identified two novel α-L-arabinofuranosidases from *B. longum* subsp. *longum* JCM 1217 that act on different linkages in arabinan. These enzymes may be required for efficient degradation and assimilation of arabinan in the probiotic bifidobacteria. The genes encoding these enzymes are located side-by-side in a gene cluster involved in metabolic pathways for plant-derived polysaccharides, which may confer adaptability in adult intestines.

## INTRODUCTION

*Bifidobacterium* spp. are gram-positive, rod-shaped, catalase-negative, strictly anaerobic bacteria. Bifidobacteria mainly inhabit the large intestines of mammals including humans. In humans, bifidobacteria are one of the dominant members of the intestinal microbiota and are believed to exert various health-promoting effects (1, 2). Bifidobacterial flora varies with age. *B. bifidum, B. breve,* and *B. longum* subsp. *infantis* are found in infant intestines and *B. adolescentis*, *B. catenulatum*, and *B. pseudocatenulatum* live in adult intestines. *B. longum* subsp. *longum* is found in intestines of both infants and adults (3). *B. longum* subsp. *longum* is also added to food products as probiotics because of its wide adaptation and beneficial health effects to the host (1).

In the large intestine where bifidobacteria reside, nutrient monosaccharides are limited. Therefore, bifidobacteria possess various glycosidases and sugar transporters to assimilate indigestible polysaccharides, oligosaccharides, and complex carbohydrates. Whole genome sequencing revealed that approximately 8.5 % open reading frames (ORFs) of *B. longum* NCC 2705 are related to glycan degradation and metabolism (4). In the past decade, several groups including ours reported that bifidobacteria in infant intestines—*B. bifidum, B. longum* subsp. *infantis,* and *B. longum* subsp. *longum*—utilize oligosaccharides from the mother’s milk (5–11) and glycoprotein glycans secreted from the gastrointestinal tract (12–15). We also reported that *B. longum* subsp. *longum* possesses a unique metabolic pathway to assimilate glycans from extensins that are plant cell-wall glycoproteins (16–18). It has been known that oligosaccharides derived from plant matrix polysaccharides, such as arabinan, arabinoxylan, and arabinogalactan, could be carbohydrate sources for adult-type bifidobacteria (19,20). However, it is poorly understood how bifidobacteria use such dietary fibers. Therefore, we investigated the mechanism of arabinan utilization in *B. longum* subsp. *longum* inhabiting the intestines of both infants and adults.

In the Carbohydrate-Active Enzymes (CAZy) database (http://www.cazy.org), there are 59 putative and experimentally characterized glycosidases belonging to 23 glycoside hydrolase (GH) families from *B. longum* subsp. *longum* JCM 1217. Of these GH families, we focused on GH43 that contains hemicellulose-degrading glycosidases including α-L-arabinofuranosidase, β-xylosidase, arabinanase, and xylanase. Nine ORFs with eleven GH43 domains occur in the genome of *B. longum* subsp. *longum* JCM 1217. Notably, five previously uncharacterized GH43 ORFs, *bllj*_1850-*bllj*_1854, form a gene cluster. In this study, we cloned and characterized two GH43 enzymes, BLLJ_1852 and BLLJ_1853, and found that these glycosidases cooperatively degrade arabinan by acting on different glycosidic bonds.

## RESULTS

### *Bifidobacterium longum* subsp. *longum* JCM 1217 utilizes arabinan as a carbon source

Sugar-beet arabinan is composed of α1,5-linked poly-L-arabinofuranose as a backbone with branched side chains of α1,2- or α1,3-linked L-arabinofuranose (21). Arabino-oligosaccharides are considered to be potential prebiotics for bifidobacteria (19,20). However, it is not known whether arabinan is utilized by *B. longum* subsp. *longum.* First, we tested whether *B. longum* subsp. *longum* JCM 1217 grows in a medium with arabinan as the sole carbon source (Fig. 1). This strain grew well in the glucose-containing medium and moderately in the L-arabinose-containing one. In the arabinan-containing medium, it reached an OD_600_ value similar to that of L-arabinose-containing medium although the growth rate was slightly slower. This result suggests that *B. longum* subsp. *longum* JCM 1217 degrades arabinan into L-arabinose and utilizes that as a carbon source.

**FIG 1.**
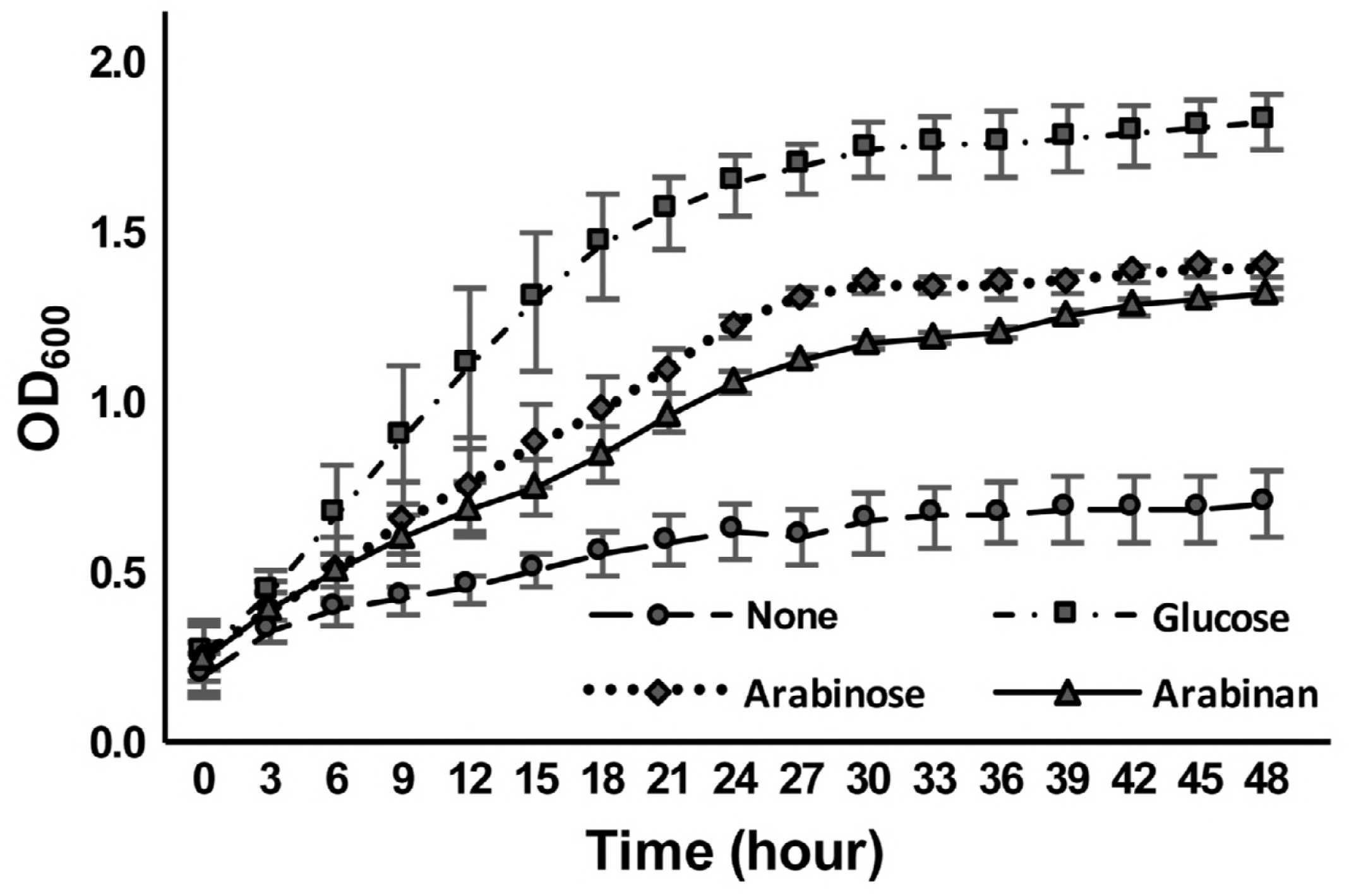
Growth of *B. longum* subsp. *longum* JCM 1217 in media containing different carbon sources. The strain was cultured at 37 °C in sugar-restricted basal medium supplemented with 1.0 % of different carbon sources under anaerobic conditions. OD_600_ was measured every 3 hours for 48 hours. Error bars indicate SD (n = 6). Symbols: circle, no sugar; triangle, arabinan; diamond, L-arabinose: square, glucose.

### Expression of *bllj*_1852 and *bllj*_1853 was enhanced in the presence of arabinan

To identify the glycosidases responsible for the degradation of arabinan, we focused on the gene cluster consisting of *bllj_1850-bllj_1854,* all of which encode GH43 domain-containing proteins. Of these, we had already determined that BLLJ_1854 was an α-L-arabinofuranosidase that preferably acted on arabinogalactan (Fujita *et al.,* submitted). Therefore, we investigated the expression of *bllj_1850-bllj_1853* in the presence of arabinan by semi-quantitative RT-PCR. The mRNA levels of *bllj_1852* and *bllj_1853* were estimated to be 4.0- and 5.4-fold higher, respectively, in arabinan medium than in glucose medium (data not shown). However, the expression of *bllj_1850* was not significantly enhanced by arabinan (less than 1.2-fold), and mRNA of *bllj_1851* could not be detected in this RT-PCR condition. This result strongly suggests that *bllj_1852* and *bllj_1853* are involved in the utilization of arabinan in this strain.

### Molecular cloning of BLLJ_1852 and BLLJ_1853

The ORF of *bllj_1852* consists of 3,741 bp and encodes a polypeptide with 1,246 residues, which is predicted to contain a signal peptide (aa 1–30), LamG domain (aa 95–253), bacterial Ig-like domain (aa 669–725), GH43 domain (aa 738–1,087), and transmembrane region (aa 1,217–1,238) (Fig. 2A). The ORF of *bllj_1853* consists of 3,294 bp and encodes a 1,097-aa polypeptide, which contains a signal peptide (aa 1–35), LamG domain (aa 81–258), GH43 domain (aa 491–782), bacterial Ig-like domain (aa 823–877), and transmembrane region (aa 1,069–1,090) (Fig. 2A). Therefore, both enzymes are predicted to be extracellular membrane-bound enzymes.

**FIG 2.**
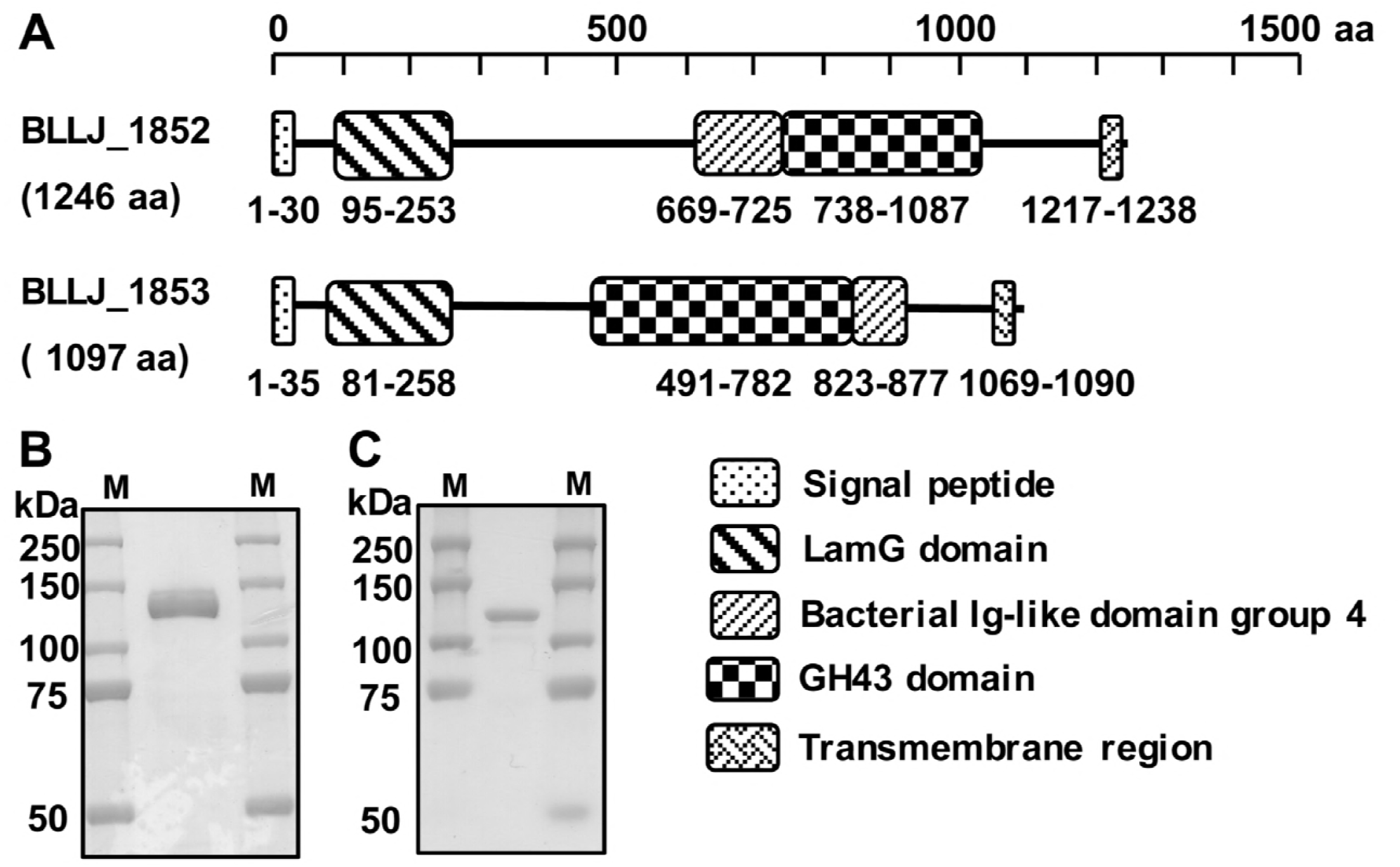
Characterization of BLLJ_1852 and BLLJ_1853. (A) Domain structure of BLLJ_1852 and BLLJ_1853. (B) SDS-PAGE of purified BLLJ_1852. (C) SDS-PAGE of purified BLLJ_1853. M, molecular size markers.

We amplified *bllj_1852* and *bllj_1853* without the regions encoding the N-terminal signal peptide and C-terminal transmembrane region using high-fidelity PCR on the genomic DNA from *B. longum* subsp. *longum* JCM 1217 as a template. The PCR products were ligated into pET23b vectors to express C-terminally 6×His-tagged proteins. *Escherichia coli* BL21(λDE3)ΔlacZ was transformed with these plasmids, and the expression was induced by adding 1.0 mM isopropyl-β-D-thiogalactopyranoside (IPTG).

The recombinant proteins were extracted using BugBuster and purified with a Ni^2+^ resin column. Purified proteins migrated as a single protein band around 130 kDa for BLLJ_1852 and 115 kDa for BLLJ_1853 on SDS-PAGE (Fig. 2B and 2C, respectively), which are consistent with the predicted molecular masses.

### Characterization of recombinant BLLJ_1852 and BLLJ_1853

The biochemical properties of recombinant BLLJ_1852 and BLLJ_1853 were investigated using pNP-α-L-arabinofuranoside (pNP-α-Ara*f*) as a substrate. The optimum pH of both glycosidases was 6.0; however, the stable pH ranges were quite different: pH 5.0–10.0 for BLLJ_1852 and pH 9.0–10.0 for BLLJ_1853 (Supplementary Fig. 1A and 1B, respectively). The optimum temperature of BLLJ_1852 and BLLJ_1853 were 45°C and 40°C, respectively. BLLJ_1852 was stable bellow 50°C for a 1 hour treatment. In contrast, BLLJ_1853 still active at 60°C for 1 hour (Supplementary Fig. 1C and 1D, respectively). Therefore, we tested the stability of BLLJ_1853 at higher temperatures and found that greater than 50 % activity was maintained below 90°C although it gradually declined (Supplementary Fig. 1E). The kinetic parameters of recombinant BLLJ_1852 for pNP-α-Ara*f* were determined: the *K*_m_, *k*_cat_, and *k*_cat_/*K*_m_ values were 9.8 mM, 19.2 s^-1^, and 1.9 s^-1^mM^-1^, respectively. These parameters of BLLJ_1853 could not be determined due to low activity for pNP-α-Araf.

### Different effects of divalent metal ions on the activities of BLLJ_1852 and BLLJ_1853

Some GH43 glycosidases were reported to have a calcium ion in their active site (22). The activities of the recombinant BLLJ_1852 and BLLJ_1853 for pNP-α-Ara*f* were measured in the presence of various divalent metal ions. BLLJ_1852 activity increased approximately 1.5-fold in the presence of all divalent metal cations tested (Zn^2+^, Mn^2^+, Cu^2+^, Ca^2+^, Co^2+^, Mg^2+^, and Ni^2+^). In contrast, BLLJ_1853 activity was not enhanced by these cations and severely inhibited by Cu^2^+ (Fig. 3A). Both glycosidases were inhibited by EDTA, suggesting a requirement for divalent metal ions (Fig. 3A). BLLJ_1852 was more sensitive to EDTA than BLLJ_1853 (Fig. 3B). Next, we tested whether these EDTA-treated enzymes could be restored or not by adding cations. BLLJ_1852 and BLLJ_1853 were treated with 100 μM EDTA at 4°C for 16 hours, and then the activities were measured in the presence of 1.0 mM CaCl_2_. The activity of BLLJ_1853 was fully restored by the addition of Ca^2^+, whereas that of BLLJ_1852 was not restored (Fig. 3C).

**FIG 3.**
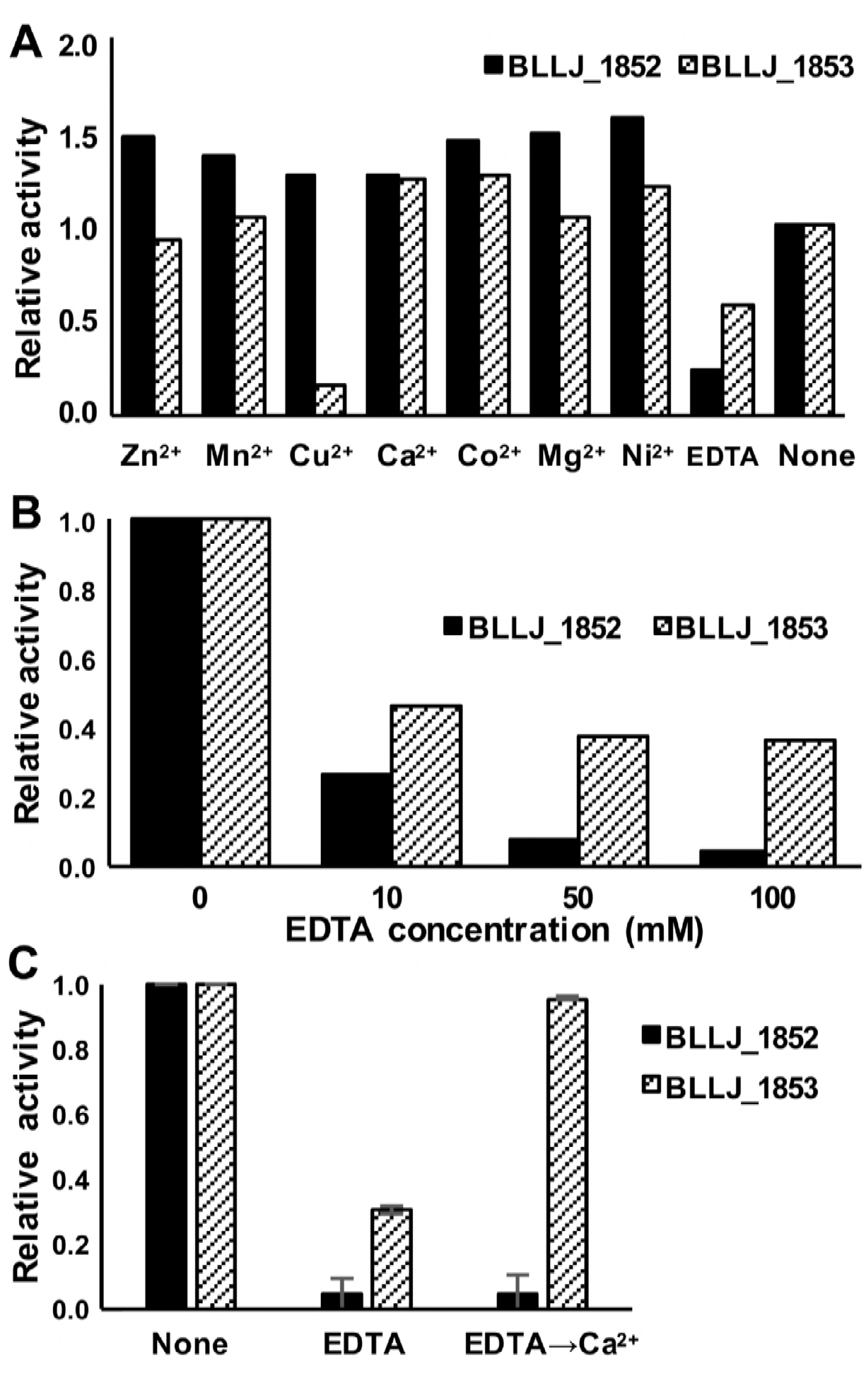
Effects of divalent metal ions on the activities of BLLJ_1852 and BLLJ_1853. (A) Effects of metal ions on the activities of recombinant enzymes. The divalent metal ions were added at 5.0 mM. (B) Effects of various concentration of EDTA on recombinant enzymes. (C) Restoration of enzyme activity by addition of Ca^2+^ after treatment with EDTA. pNP-α-Ara*f* was used as a substrate in all experiments. Error bars indicate SD (n=3).

### Substrate specificities of BLLJ_1852 and BLLJ_1853

As described above, both enzymes hydrolyzed pNP-α-Ara*f*, indicating that they are exo-α-L-arabinofuranosidases. We tested the activities of these enzymes for other pNP-glycosides, such as pNP-α-L-arabinopyranoside, pNP-β-xylopyranoside, and pNP-β-galactopyranoside, but no hydrolysis was observed (Fig. 4A). Therefore, these glycosidases have a strict specificity for α-linked L-arabinofuranose. Next, we tested substrate specificities for natural substrates. Of the natural substrates tested, BLLJ_1852 released L-arabinose from arabinan and arabinoxylan, and BLLJ_1853 released L-arabinose from arabinan only (Figs. 4B and 4C, respectively). There are α1,2- and α1,3-arabinofuranosyl side-chins on the α1,5-poly-L-arabinofuranosyl backbone of arabinan. The same α1,2- and/or α1,3-arabinofuranosyl side-chins are on the xylan backbone of arabinoxylan. Next, we tested linkage specificities using synthetic arabinobioses: Ara*f*α1,2Ara*f*α-OMe, Ara*f*α1,3Ara*f*α-OMe, and Ara*f*α1,5Ara*f*α-OMe. BLLJ_1852 acted on the α1,2- and α1,3-linkages, whereas BLLJ_1853 acted only on the α1,5-linkage (Fig. 4D). These results strongly suggest that BLLJ_1852 hydrolyzes the side chains of arabinan and arabinoxylan, and BLLJ_1853 hydrolyses the backbone of arabinan. Finally, we quantified the release rate of L-arabinose from five substrates (Table 1). Both enzymes showed the highest activity with arabinan. BLLJ_1852 also acted on arabinoxylan, but the rate was only 0.6 % of that for arabinan. Synthetic glycosides, pNP-α-Ara*f* and 4-methylumbelliferyl-α-Ara*f* (4MU-α-Ara*f*), were very slowly hydrolyzed by both enzymes. These results indicate that both enzymes have strict aglycone and linkage specificities.

**FIG 4.**
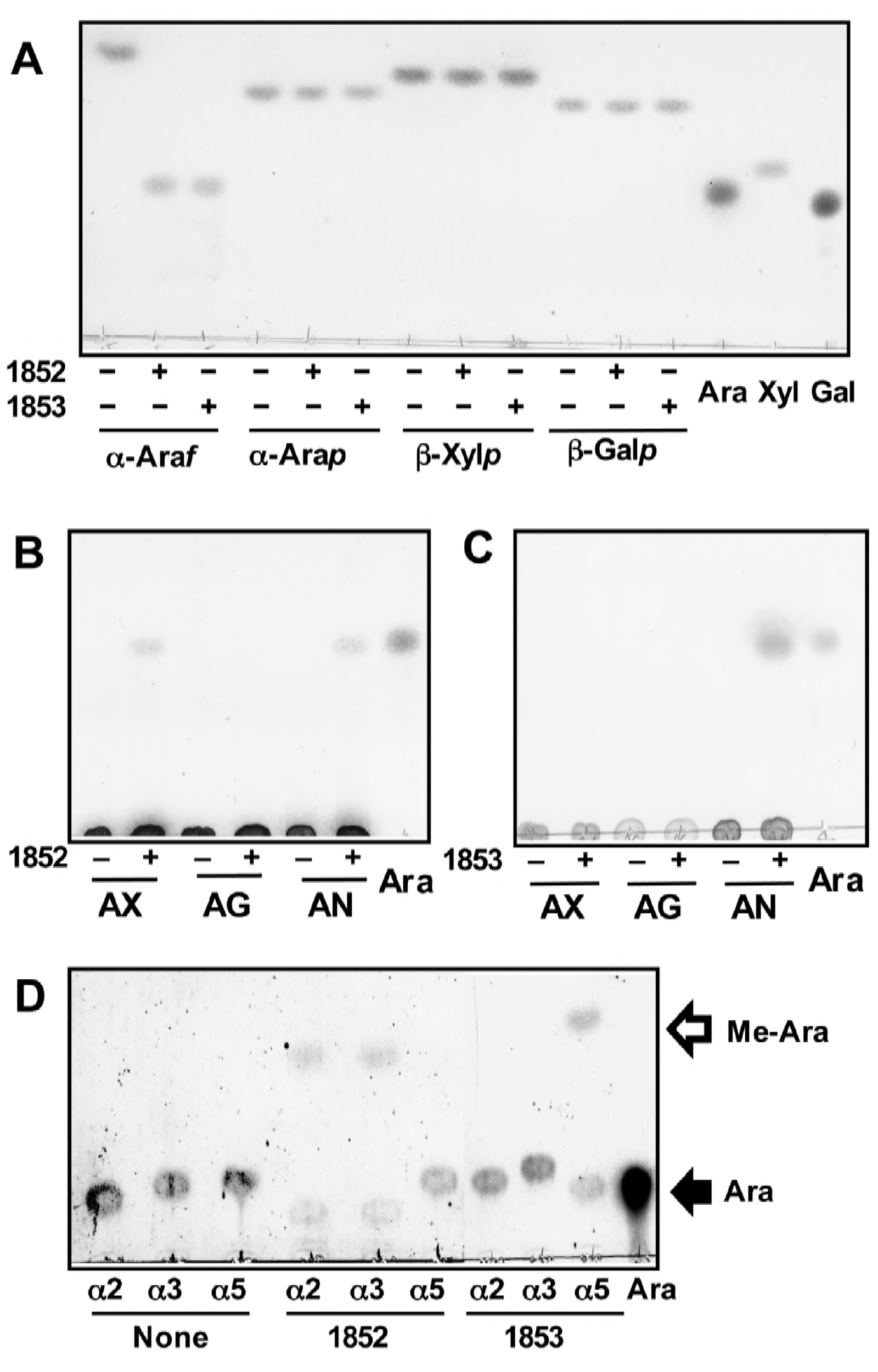
Substrate specificities of BLLJ_1852 and BLLJ_1853. The substrates were incubated with recombinant BLLJ_1852 or BLLJ_1853 at 37°C overnight and analyzed by TLC. (A) Action on synthetic substrates. α-Ara*f* pNP-α-L-arabinofuranoside; α-Arap, pNP-α-L-arabinopyranoside; β-Xyl*p*, pNP-β-xylopyranoside; β-Galp, pNP-β-galactopyranoside; Ara, standard L-arabinose; Xyl, standard xylose; Gal, standard galactose. (B) Action of BLLJ_1852 on natural substrates. AX, arabinoxylan; AG, arabinogalactan; AN, arabinan; Ara, standard L-arabinose. (C) Action of BLLJ_1853 on natural substrates. AX, arabinoxylan; AG, arabinogalactan; AN, arabinan; Ara, standard L-arabinose. (D) Action of BLLJ_1852 and BLLJ_1853 on synthetic disaccharide substrates. α2, Ara*f*α1,2Ara*f*α-OMe; α3, Ara*f*α1,3Ara*f*α-OMe; α5, Ara*f*α1,5Ara*f*α-OMe; Ara, standard L-arabinose. The developing solvent used in panels A-C was 1-Butanol:acetic acid:water (2:1:1, by volume); chloroform:methanol:acetic acid (6:1:1, by volume) was used for panel D.

**TABLE 1.** Specific activities of BLLJ_1852 and BLLJ_1853 for various substrates Reaction mixtures were incubated at 37°C for 30 min. The released L-arabinose was determined by the enzymatic measurement method, and the relative values are shown. pNP, pNP-α-L-arabinofuranoside; 4MU, 4-methylumbelliferyl-α-L-arabinofuranoside; AN, sugar-beet arabinan; AX, wheat arabinoxylan; AG, larch wood arabinogalactan; nd, not detected.

|  | pNP | 4MU | AN | AX | AG |
| --- | --- | --- | --- | --- | --- |
| BLLJ_1852 | 1.00 | 0.62 | 70.23 | 0.62 | nd |
| <u>BLLJ_1853</u> | <u>0.009</u> | <u>0.23</u> | <u>4.42</u> | <u>nd</u> | <u>nd</u> |

Reaction mixtures were incubated at 37°C for 30 min. The released L-arabinose was determined by the enzymatic measurement method, and the relative values are shown. pNP, pNP-α-L-arabinofuranoside; 4MU, 4-methylumbelliferyl-α-L-arabinofuranoside; AN, sugar-beet arabinan; AX, wheat arabinoxylan; AG, larch wood arabinogalactan; nd, not detected.

## DISCUSSION

GH43 is a large family composed of more than 10,000 sequences and currently classified into 37 subfamilies (23). Nine enzymes with eleven GH43 domains from *B. longum* subsp. *longum* JCM 1217 are listed in the CAZy database. They are highly diversified with ten domains belonging to eight different subfamilies and one remaining unclassified. Four GH43 enzymes, including BLLJ_1852 and BLLJ_1853, have been characterized. BLLJ_0213, belonging to subfamily 29 of GH43 (GH43_29), is a HypAA α-L-arabinofuranosidase specific for α1,3-linked Ara*f* in Ara*f*α1,3Ara*f*β1–2Ara*f*β1-hydroxyprolin of plant glycoproteins (24). BLLJ_1840, belonging to GH43_24, is an exo-β1,3-galactanase that acts on the backbone of type II arabinogalactan (25). In this study, we found that BLLJ_1852 (GH43_27) and BLLJ_1853 (GH43_22), which are exo-α-L-arabinofuranosidases, cooperatively degrade arabinan. BLLJ_1852 hydrolyzed Ara*f*α1,2Ara*f*α-OMe and Ara*f*α1,3Ara*f*α-OMe, suggesting that it removes α1,2- and α1,3-linked Ara*f* side chains from arabinan. BLLJ_1853 specifically hydrolyzed Ara*f*α1,5Ara*f*α-OMe, indicating that it processively degrades α1,5-linked L-arabinose of the arabinan backbone from the non-reducing terminus. BLLJ_1852 also acted very slowly on arabinoxylan, which contains Ara*f*α1,2/3Xyl*p*β1-R, but not on arabinogalactan, which contains Ara*f*α1,3Gal*p*β1-R. Since Ara*f* Xyķ, and Galp have equatorial hydroxyl groups at the C2 and C3 positions but only Galp has an axial hydroxyl group at the C4 position, BLLJ_1852 may have a strict recognition of equatorial hydroxyl groups at the C4 position in aglycone.

Our preliminary results suggest that BLLJ_1850 (GH43_22_34), BLLJ_1851 (GH43_unclassified subfamily_26), and BLLJ_1854 (GH43_22) are also α-L-arabinofuranosidases. BLLJ_1850 rapidly acts on arabinoxylan and weakly on arabinogalactan, and BLLJ_1851 rapidly acts on arabinan and weakly on arabinoxylan. The details will be reported elsewhere (Komeno *et al*., in preparation). BLLJ_1854 rapidly acts on arabinogalactan and weakly on arabinan (Fujita *et al.,* submitted). Except for BLLJ_0213, these GH43 enzymes form a gene cluster, and there is almost no functional redundancy (Fig. 5A).

**FIG 5.**
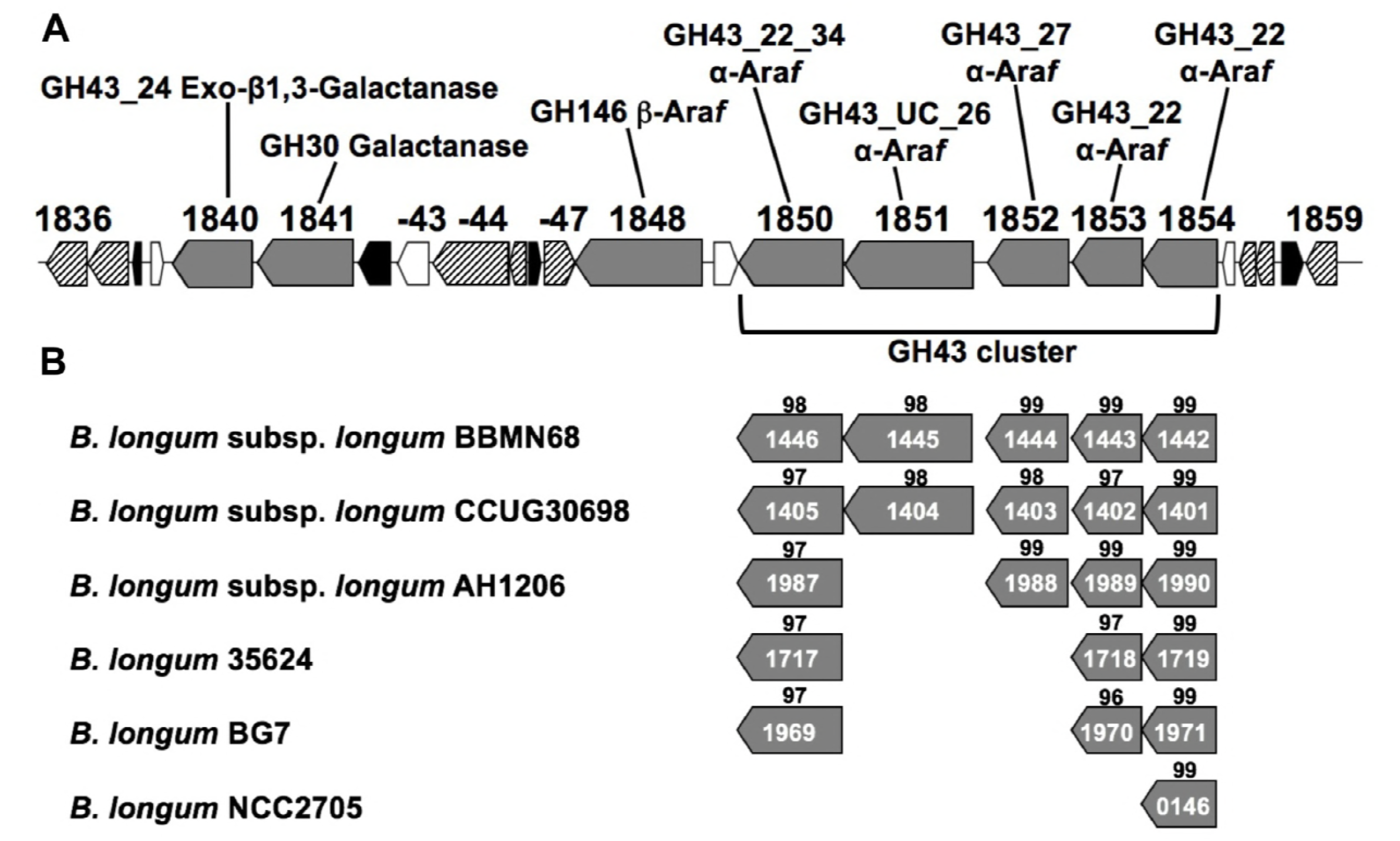
(A) Map of the hemicellulose-degrading gene cluster in *B. longum* subsp. *longum* JCM 1217. Gray boxes, glycosidases; striped boxes, sugar transporters; black boxes, transcriptional regulators; white boxes, hypothetical proteins. BLLJ_1850 and BLLJ_1850 have two GH43 domains, and the N-terminal GH43 domain of BLLJ_1850 belongs to an unclassified (UC) subfamily. (B) Conservation of the GH43 cluster in various *B. longum* strains. Scores indicated above the boxes indicate percentage identity of amino acid sequences with those of *B. longum* subsp. *longum* JCM 1217.

GH43_27 is a small subfamily whose members are restricted in bacterial enzymes. So far only Abf43B α-L-arabinofuranosidase from *Paenibacillus* sp. E18 (GenBank accession number AFC38437) (26) and GbtXyl43B β-D-xylosidase from *Geobacillus thermoleovorans* IT-08 (ABD48561) (27) have been experimentally characterized. The former hydrolyzed pKP-α-L-Ara*f* but neither arabinan nor arabinoxylan, and the latter hydrolyzed pNP-β-D-Xyķ and xylan. Since BLLJ_1852 is an arabinan-degrading exo-1,2–1,3-α-L-arabinofuranosidase, the substrate specificity of BLLJ_1852 is quite different from those of Abf43B and GbtXyl43B. The amino acid sequence of the GH43_27 domain (aa 738–1,087) of BLLJ_1852 had only 10 and 9 % identity with the corresponding domains of Abf43B and GbtXyl43B, respectively. Adult-type bifidobacteria have putative homologs with higher identities of 45–50 %: BAD_0149 from *B. adolescentis* ATCC 15703, BBCT_0647 from *B. catenulatum,* BBPC_0158 from *B. pseudocatenulatum,* and BBSC_0258 from *B. scardovii.* However, within the closely related strains of *B. longum* whose genomic data are available, the BLLJ_1852 homolog was lost in several strains, such as *B. longum* 35264, *B. longum* BG7, and *B. longum* NCC2705 (Fig. 5B), and is also missing in *B. longum* subsp. *infantis.*

In comparison with GH43_27, the subfamily GH43_22 is relatively larger and more widely distributed in eukaryote (several fungi) and archaea in addition to bacteria. In this subfamily, we could not find an experimentally characterized enzyme. In our strain, there are three enzymes belonging to this subfamily, BLLJ_1850 (GH43_22_34), BLLJ_1853, and BLLJ_1854, which exhibit distinct substrate preference for arabinoxylan, arabinan, and arabinogalactan, respectively, as described above. Within *B. longum* strains, the homologs of BLLJ_1853 and BLLJ_1850 are conserved expect for *B. longum* NCC2705, and those of BLLJ_1854 are conserved in all strains (Fig. 5B). BLLJ_1853 is arabinan-degrading exo-1,5-α-L-arabinofuranosidase. The enzymes with similar substrate specificity are classified in the subfamily GH43_26: Araf43A (BAL68753) from *Streptomyces avermitilis* MA-4680 (28, 29), and Abf43K (ACE82749) and Abf43L (ACE84379) from *Cellvibrio japonics* Ueda107 (30). However, BLLJ_1853 shares very low identities with the amino acid sequences of Araf43A (13%), Abf43K (11%), and Abf43L (15%), suggesting that the evolutionary origin may be different. The exo-1,5-α-L-arabinofuranosidase that belongs to the other families of GH51 was also reported: AbfA (CAA99595) from *Bacillus subtilis* strain 168 (31).

It is known that *B. longum* subsp. *longum* is widely distributed in the intestines of infants, adults, and elderly hosts. A recent study on the comparative genome analyses of human isolates revealed that this GH43 gene cluster is enriched in strains from elderly subjects (3). These findings suggest that this gene cluster is generally important for the colonization of *B. longum* subsp. *longum* in the intestines of adults who consume vegetables and grains.

## MATERIALS AND METHODS

### Bacterial strains and culture

*B. longum* subsp. *longum* JCM 1217 was obtained from the Japan Collection of Microorganisms (RIKEN Bioresource Center, Japan). The strain was cultured in GAM broth (Nissui Pharmaceutical, Japan) or sugar-restricted basal medium (1.0 % Bacto peptone, 0.5 % Bacto yeast extract, 0.5 % sodium acetate trihydrate, 0.2 % di-ammonium hydrogen citrate, 0.08 % L-cysteine hydrochloride monohydrate, 0.02 % magnesium sulfate heptahydrate, 1.36 % L-ascorbic acid, and 0.44 % sodium carbonate anhydrous) at 37°C under anaerobic conditions using the AnaeroPack Anaero (Mitsubishi Chemical, Japan). *Escherichia coli* DH5α and *E. coli* BL21(λDE3)Δ*lac*Z (a gift from Prof. T. Katayama, Kyoto University, Kyoto, Japan) were cultured in Luria-Bertani (LB) broth (BD, USA) at 37°C under aerobic conditions.

### Chemicals and substrates

*p*-Nitrophenyl (pNP)-glycosides were purchased from Sigma-Aldrich, MO, USA; sugar-beet arabinan and wheat arabinoxylan were purchased from Megazyme, Ireland; larch wood arabinogalactan was purchased from Tokyo Chemical Industry, Japan; and L-arabinose was purchased from Wako Chemicals, Japan. Synthetic 1-*O*-methylated arabinobioses (Ara*f*α1,2Ara*f*α-OMe, Ara*f*α1,3Ara*f*α-OMe, and Ara*f*α1,5Ara*f*α-OMe) were gifted from Prof. S. Kaneko, Ryukyu University, Japan (32). Other chemicals were obtained from Nacalai Tesque Inc., Japan.

### Growth assay of *B. longum* subsp. *longum*

*B. longum* subsp. *longum* JCM 1217 was inoculated in sugar-restricted basal medium supplemented with 1.0 % L-arabinose and cultured at 37°C under anaerobic conditions until the optical density at 600 nm (OD_600_) reached 1.0. A 10 % volume seed culture was added to basal medium containing 2.0 % arabinan, 2.0 % L-arabinose, or 2.0 % glucose. The wells of a 96-well plate were filled with these mixtures followed by sealing with Microseal B (Bio-Rad, USA). The plate was maintained at 37°C, and the OD_600_ was monitored every 3 hours using Powerscan HT (Dainippon Sumitomo Pharma, Japan).

### Semi-quantitative RT-PCR

*B. longum* subsp. *longum* JCM 1217 was cultured at 37°C under anaerobic conditions in sugar-restricted basal medium with either 2.0 % arabinan or 2.0 % glucose. RNA extraction was carried out using the TRIzol Reagent (Thermo Fisher Scientific, MA, USA) and purification used the PureLink RNA Mini Kit (Thermo Fisher Scientific). Reverse transcription was performed using PrimeScript Reverse Transcriptase (Takara Bio Inc., Japan). Primers for RT-PCR were designed as follows: 5’-CCCAAGCTTGATACCACCGATTCATCGGCCGC and 5’-GTGCCGCTCGAGGGAGATGACGGCACCCGGCTTCTTG for *bllj_1850,* 5’-CGCGGATCCGGACGACGCGACGCCTGCGGTG and 5’-CCCAAGCTTAGAGGACGGCAGACCGGAGTCTGC for *bllj_1851,* 5’-GGAATTCCATATGGATACCGTTCCGACCAATAATCTCATC and 5’-CCCAAGCTTAGACAGACCGAGCTTGTTGCCCG for *bllj_1852,* and 5’-CGGAATTCGGAGAGCGCATCGCCAATCGATG and 5’-GTGCCGCTCGAGGGTGTTGGACAGAGCGCTGCCCGG for *bllj_1853.* RT-PCR was performed using TaKaRa Ex Taq HS (Takara Bio Inc.). PCR products were separated by electrophoresis on a 0.8 % agarose gel. The gel staining used SYBR Gold Nucleic Acid Gel Stain (Thermo Fisher Scientific). The densities of the bands were quantified using the ImageJ program.

### Cloning, expression, and purification of BLLJ_1852 and BLLJ_1853

The DNA fragment for base pairs 106–3645 of *bllj_1852* (GenBank accession number BAJ67516) without the coding sequences of the N-terminal signal peptide and C-terminal transmembrane region was amplified by high-fidelity PCR using KOD plus ver. 2 (Toyobo, Japan), genomic DNA from *B. longum* subsp. *longum* JCM 1217 as a template. The same pair of primers for RT-PCR with the NdeI and HindIII sites were used. The amplified fragment was ligated into the NdeI and HindIII sites of the pET23b(+) vector (Novagen, USA). Similarly, the DNA fragment of base pairs 91–3203 of *bllj_1853* (BAJ67517) was amplified using same the pair of primers as the RT-PCR containing the EcoRI and XhoI sites. The product was ligated into corresponding sites of pET23b(+). The nucleotide sequences were confirmed by sequencing. The *E. coli* BL21(λDE3)ΔlacZ strain was transformed with these plasmids and cultured in LB liquid medium containing 100 μg/mL ampicillin at 37°C under aerobic conditions until the OD_600_ reached 0.35. Then the flask was put on ice in order to lower the temperature. Next, to induce expression, 0.1 mM isopropyl-β-D-thiogalactopyranoside (IPTG) was added to the culture, which was further incubated at 25°C for 15 hours. The cells were harvested by centrifugation and lysed with the BugBuster Protein Extraction Reagent (Novagen). The cell free protein extract was applied to a His GraviTrap (GE Healthcare, USA), and adsorbed proteins were eluted by a stepwise imidazole concentration gradient in 20 mM sodium acetate buffer at pH 6.5 containing 500 mM NaCl. The active fraction was dialyzed with a size 36 dialysis membrane (Wako Chemicals) against 0.5 mM sodium acetate buffer at pH 6.5 overnight at 4°C.

Protein concentration was measured using a Bio-Rad Protein Assay Dye Reagent Concentrate (Bio-Rad).

### Glycosidase assay

α-L-Arabinofuranosidase activity was measured based on the release of pNP from pNP-α-Araf The reaction mixture containing 1.0 mM pNP-Ara*f*, an appropriate amount of purified enzyme, and 1.0 mM sodium acetate buffer at pH 5.5 to a total volume of 50 μL was incubated at 37°C. After the incubation, 1.5 volumes of 1.0 M Na_2_CO_3_ was added as a stop solution, and the released pNP was measured at 405 nm. For optimum pH tests, sodium acetate (pH 3.0–6.0), sodium phosphate (pH 6.5–7.5), and Tris-HCl buffers (pH 8.0–10.0) were used. The 50 μL reaction mixture containing 1.0 mM of each pH buffer, 1.0 mM pNP-α-Ara*f*, and 5 μg enzyme was incubated at 37°C for 30 min (BLLJ_1852) or overnight (BLLJ_1853). For a stable pH assay, the enzyme was pretreated in 10 mM buffer at 4°C for 24 hours, and then the remaining activity was measured in sodium acetate buffer at pH 5.5. For the metal ion test, 5 mM chloride salts (ZnCl_2_, MnCl_2_, CuCl_2_, CaCl_2_, CoCl_2_, MgCl_2_, and NiCl_2_) or EDTA 2Na were added to reaction mixtures. For the assays for natural substrates, such as arabinan, arabinoxylan, and arabinogalactan, the reaction mixture containing 0.5 % substrate, 1.0 mM sodium acetate buffer at pH 5.5, and the indicated amount of enzyme was incubated at 37°C for the appropriate period. The reaction mixture was analyzed by TLC or applied to enzymatic measurements of released L-arabinose using an L-arabinose/D-galactose assay kit (Megazyme).

### TLC

TLC analysis was carried out using silica gel 60 plates (Merck, Germany) with 1-butanol:acetic acid:water (2:1:1 by volume) or chloroform:methanol:acetic acid (6:1:1 by volume) as developing solvents. Sugars were visualized by spraying acetone:aniline:diphenyl amine:phosphoric acid (100:1:1:10 by volume) and heating to 150°C for 15 min.

## SUPPLEMENTAL MATERIAL

Supplemental material for this article may be found at *###*

SUPPLEMENTAL FILE 1, PDF file, ## MB.

## ACKNOWLEDGMENTS

We thank Prof. S. Kaneko (University of Ryukyus, Japan) for the kind gifts of Ara*f*α1,2Ara*f*α-OMe, Ara*f*α1,3Ara*f*α-OMe, and Ara*f*α1,5Ara*f*α-OMe and Prof. T. Katayama (Kyoto University, Japan) for providing *E. coli* BL21(λDE3)Δ*lacZ*. This work was supported by JSPS KAKENHI grants 15K07448 and 18K05494 (to H.A.).

We declare that there is no conflict of interest.

---

**Fig. S1** General properties of recombinant BLLJ_1852 and BLLJ_1853. pNE-α-Ara*f* was incubated with recombinant enzymes at 37°C. (A) Effect of pH on BLLJ_1852. (B) Effect of pH on BLLJ_1853. (C) Effect of temperature on BLLJ_1852. (D) Effect of temperature on BLLJ_1853. (E) Thermal stability of BLLJ_1853 at high temperatures. Error bars indicate SD (n=3). For stability tests, enzymes were pre-incubated in each pH buffer for 16 hours at 37°C or in sodium acetate buffer (pH 6.0) at each temperature for 1 hour.

